# Global & Temporal Patterns of Submicroscopic *Plasmodium falciparum* Malaria Infection

**DOI:** 10.1101/554311

**Authors:** Charles Whittaker, Hannah Slater, Teun Bousema, Chris Drakeley, Azra Ghani, Lucy Okell

**Affiliations:** MRC Centre for Global Infectious Disease Analysis, Department of Infectious Disease Epidemiology, Imperial College London, London, United Kingdom; PATH, 2201 Westlake Avenue, Seattle, USA; Department of Medical Microbiology, Radboud University Medical Center, Nijmegen, The Netherlands; Department of Immunology and Infection, Faculty of Infectious and Tropical Diseases, London School of Hygiene and Tropical Medicine, London, United Kingdom

## Abstract

Molecular detection of *Plasmodium falciparum* infection has revealed large numbers of individuals with low-density (yet transmissible) infections undetectable by microscopy. Here we present an updated systematic review of cross-sectional malaria surveys to explore the prevalence and drivers of these submicroscopic infections and define where they are likely to be relevant to malaria control efforts. Our results show that submicroscopic infections predominate in low transmission settings, but also reveal marked geographical variation in their prevalence, being highest in South American surveys and lowest in West African studies. Whilst current transmission levels partly explain these results, we find that historical transmission intensity also represents a crucial determinant of the size of the submicroscopic reservoir. Submicroscopic infection was more likely in adults than children, although we did not observe a statistically significant influence of seasonality. Our results suggest that the contribution of submicroscopic infections to transmission likely varies substantially across settings, potentially warranting different approaches to their targeting in the approach to elimination.

## Introduction

The ability to accurately detect malaria infection during population surveys in endemic areas is a cornerstone of effective surveillance and control of the parasite. Routinely, this is undertaken using microscopy of blood films or rapid diagnostic tests, although recent years have seen increased usage of more sensitive molecular methods in research contexts. These techniques (typically PCR-based^1^) have revealed the widespread presence of infections with parasite densities lower than the threshold of detection by routine methods^2–8^.

Such “submicroscopic” infections are present across a range of different settings and populations^5,9–11^. Although rarely causative of severe symptoms, they have been associated with a number of adverse outcomes during pregnancy^12,13^, as well as mild anaemia^14^ and other symptoms (vomiting, jaundice etc.) in children under 10^15^. In addition to their capacity to cause low-grade disease, these infections may hold public health relevance due to their contribution to onwards transmission. Although typically characterised by lower parasite densities than microscopically detectable infections, individuals with submicroscopic infections frequently harbour gametocytes (the transmissible form of the parasite) and are capable of contributing to onwards transmission^16,17^. Recent work suggests that there is often overlap in the parasite densities (as assessed by quantitative PCR) of individuals that are detectable by microscopy and those who are submicroscopically infected, suggesting that differences in parasite densities (and by extension, infectivity) between these infections may not always be that significant in practice^18^. Submicrosopically infected individuals have been shown to contribute to transmission of the parasite across both areas of high^19^ and low^20^ transmission intensity, as well seasonal^16^ and perennial^17^ settings, further underscoring the potential relevance of this infected sub-group to malaria control efforts.

Despite this, our understanding of the factors influencing the size of the submicroscopic reservoir across endemic populations remains far from complete. Previous systematic reviews have found that microscopy misses on average, half of all *P. falciparum* infections compared to PCR-based methods in endemic country cross-sectional surveys^21^ and that adults are more likely to harbour submicroscopic infections than children^22^. However, these reviews also identified extensive unexplained variation in the size of the submicroscopically infected population across settings, suggesting the existence of other important factors that determine the size of the submicroscopic reservoir. For example, although the burden of submicroscopic infection is highly heterogeneous across different locations^9,23^, it remains unclear whether this represents systematic variation according to geography or is reflective of other underlying location-specific characteristics.

Resolving these gaps in our understanding of submicroscopic epidemiology has material consequences for the future of malaria control. Despite mixed reports surrounding more recent progress^24^, transmission is still declining in many endemic countries and the WHO has identified 21 countries that have the potential to eliminate by 2020 (http://www.who.int/malaria/areas/elimination/e2020/en/). Low transmission settings such as these can possess high proportions of submicroscopically infected individuals^25,26^: the existence of a significant submicroscopic infection prevalence in an otherwise asymptomatic, microscopically negative population in Ethiopia^27^ highlights that malaria infections can persist in populations entirely absent of clinical indicators or detectable using typically deployed diagnostics. Understanding the prevalence, detectability and infectiousness of low-density infections in these settings will be essential for planning for elimination: is there benefit in detecting and treating such infections, or are resources better spent elsewhere? Improving our understanding of the drivers of submicroscopic infection in these low transmission areas is therefore crucial to enable better definition of when and where submicroscopic infection is likely to occur and how the size of the submicroscopic reservoir is likely to change as areas approach elimination.

Here we update previous reviews on submicroscopic malaria infection prevalence^21,25^, leveraging the increase in the usage of molecular methods over the past 5 years to explore novel determinants of submicroscopic infection prevalence. These include geographical location and historical patterns of transmission, seasonality and the role of age at a finer resolution than previously possible. These results are then integrated with literature-based estimates of the infectivity of submicroscopic individuals to mosquitoes in order to estimate their contribution to malaria transmission across a range of different settings.

## Results

### Microscopic Detectability of Malaria Infection Across the Endemicity Spectrum

Across the 267 surveys included in our analyses that assessed infection status across a range of ages, microscopy detected on average 46.5% of all PCR-detectable infections (43.0% - 50.0%, 95% CI), although this varied substantially across settings **(Fig.1A)**. There was no difference in the observed relationship between PCR and microscopy prevalence when the model was fitted separately to data collated in previous reviews and the data newly extracted here **(Supp Fig.2 and Supp Fig.3)**: A number of more flexible model structures were also fitted to the data, although a log-linear model provided the best overall fit (**Supp Fig.4**). The overall fitted relationship between PCR prevalence and microscopy prevalence was:

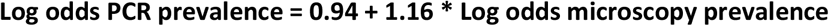

where log odds = ln(prevalence/(1-prevalence) and prevalence = exp(log odds)/(1+exp(log odds)). The prevalence ratio (defined as the proportion of PCR positive infections also detectable by light microscopy) increased as malaria transmission (measured by Survey PCR Prevalence) increased, indicating a declining proportion of submicrosopically infected individuals, from between 70-80% in the areas of lowest PCR prevalence to only 10-20% in the highest prevalence areas **(Fig.1B)**. However, there remained large variation in the proportion of submicroscopic infections between surveys carried out in areas with similar PCR prevalence. Grouping surveys by global region (West Africa, East Africa, South America, and Asia/Oceania) revealed marked geographical variation in the prevalence ratio (ANOVA, p<0.001, df=3), being lower in South American surveys compared to other regions (Tukey’s HSD, p<0.001 for all 3 pairwise comparisons) **(Fig.1C).** There was no statistically significant effect of sampling season after controlling for survey PCR prevalence (ANOVA, df=2, p=0.181, **Supp Fig.5**), but a statistically significant effect of PCR methodology (ANOVA, df=5, p<0.001, **Supp Fig.6**). Scanning a higher number of microscopy fields to determine infection presence/absence was also significantly associated with the prevalence ratio increasing (ANOVA, df=1, p<0.01).

**Figure 1:**
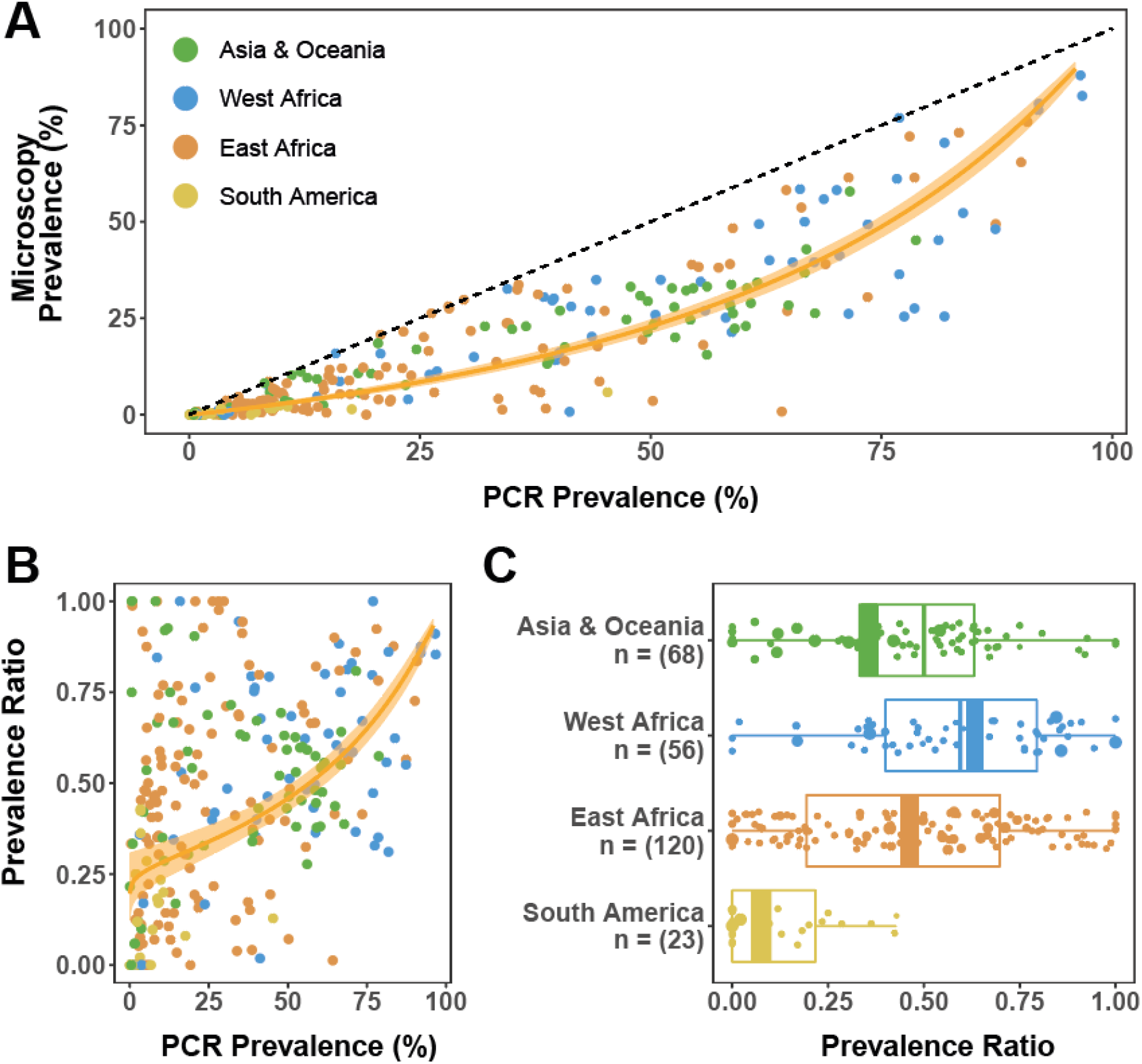
Prevalence of infection by PCR versus microscopy in 267 prevalence survey pairs and model fits. Bayesian Markov chain Monte Carlo methods were used to fit a linear relationship between PCR prevalence and microscopy prevalence on the log odds scale. **(A)** Microscopy and PCR prevalence (n = 267) in surveys identified through this and previous systematic reviews. The fitted model relationship (yellow line) and the 95% credible interval of the mean (paler shaded area). **(B)** The prevalence ratio (the proportion of PCR positive individuals also detectable by microscopy) according to underlying PCR prevalence for each of the 267 survey microscopy-PCR pairs (points) used to fit the full model. The estimated average prevalence ratio (yellow line) and 95% credible interval of the mean (paler shaded area) are also shown. **(C)** Histogram of the prevalence ratio disaggregated by global region. For each region, the size of the point reflects the number of individuals tested by microscopy and PCR. Thick coloured bar on the histogram represents the weighted mean prevalence ratio for each global region.

### Global Variation in the Size of the Submicroscopic Reservoir

Average transmission intensity (a factor that significantly influences the prevalence ratio, measured here by PCR prevalence) differed significantly between the 4 global regions. It was therefore unclear whether the observed differences in the prevalence ratio were directly related to geography, or whether these results were confounded by the differences in average transmission intensity across the global regions. To account for this, we fitted separate log-linear models to the data from each global region and assessed the modelled prevalence ratio across the range of transmission intensities found in each global region **(Fig.2A-2D)**. These results revealed that the prevalence ratio in surveys from South America was lower than would be expected based on their respective transmission intensities alone, and consistently lower than all other global regions **(Fig.2E).** This points to the role of additional factors in shaping the relationship between transmission and microscopy sensitivity other than just endemicity levels. Importantly, there was no evidence that these differences were due to PCR and microscopy methodologies used in different global regions, with no substantial variation in the methodology used between regions **(Supp Fig.7).** This highlights that the low prevalence ratio in South America (indicating a wider disparity in PCR and microscopy performance) was likely not occurring due to systematic differences in assay sensitivities between settings.

**Figure 2:**
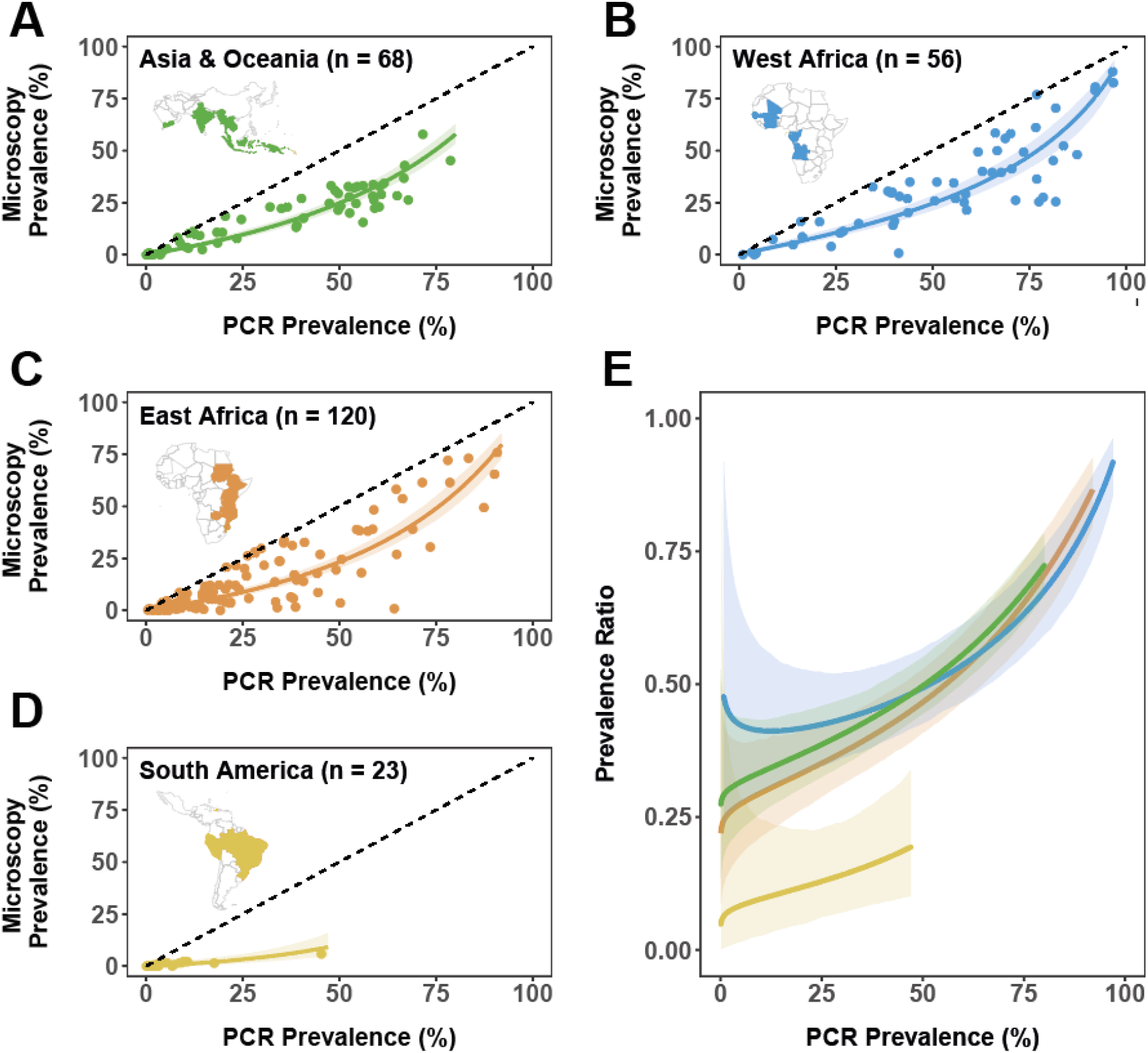
Global variation in the prevalence ratio and the relative size of the submicroscopic reservoir. Microscopy and PCR prevalence in included surveys (points), the model-fitted relationship (coloured line) and 95% credible interval (shaded area) for **(A)** West Africa **(n = 56), (B)** East Africa **(n = 120), (D)** South America **(n = 23)** and **(E)** Asia and Oceania **(n = 66). (C)** The model-fitted average microscopy: PCR prevalence ratio by PCR prevalence for each of the 4 regions (coloured) line and 95% credible interval (shaded area). Coloured countries on each regional map indicates countries for which studies were identified during the systematic review.

### Exploring the Influence of Historical and Current Transmission Patterns

The majority of the South American surveys had been undertaken in areas marked by historically low transmission. We therefore investigated whether a high proportion of submicroscopic infections (low prevalence ratio) might be observed in areas around the world that had experienced similar historically low patterns of transmission. Each survey carried out in Africa (a continent for which estimates of malaria prevalence are available at a high spatio-temporal resolution) was geolocated and the estimates of Historical and Current Regional Prevalence at the administrative unit 1 level collated to disentangle the effects of historical prevalence from more contemporary transmission. Note that we distinguish between local malaria transmission (defined by the prevalence recorded in each survey, hereafter referred to as Survey PCR Prevalence), and malaria transmission at the regional level (reflecting broader patterns of transmission). This regional-level transmission represents the average of a heterogeneous mixture of higher and lower transmission areas, and has relevance to local transmission because factors like human movement patterns and circulating parasite genetic diversity are often similar across nearby settings in the same region, even if transmission levels markedly differ^28,29^.

Our results indicated that Regional Historical Prevalence, but not Regional Current Prevalence was a significant predictor of the prevalence ratio when controlling for Survey PCR Prevalence (ANOVA, p<0.001 for both Survey Prevalence and Regional Historical Prevalence, p=0.73 for Regional Current Prevalence), suggesting that historical patterns of transmission are an important determinant of the submicroscopic reservoir size. To further explore this, we classified each survey in our review from Africa into 3 transmission “archetypes” based on historical and current regional levels of transmission and fitted regression models to each. The results from these models were concordant with the results from the ANOVA, with African surveys in regions with both historically and currently low transmission (Sudan, Ethiopia and parts of Kenya and Tanzania) having on average, a lower prevalence ratio (more submicroscopic infections) compared to other low-endemic areas in Africa where historical transmission has been high **(Fig.3B and Fig.3C)**. There was no evidence of systematic differences in the PCR and microscopy methodologies in different transmission archetypes **(Supp Fig.8)** and the observed results were robust to the choice of High/Low stratification threshold **(Supp Fig.9)**.

**Figure 3:**
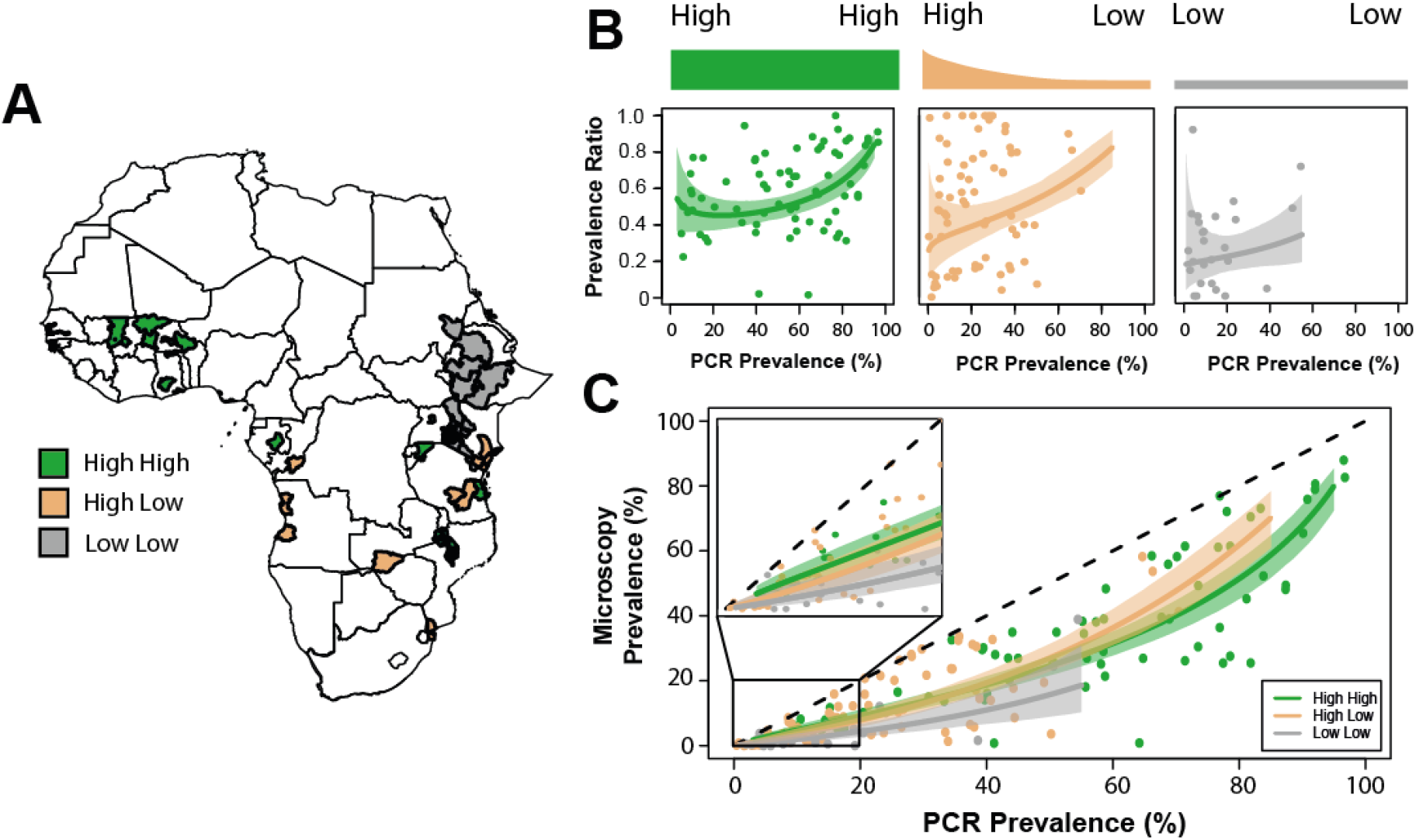
The effect of historical and current transmission intensity on the prevalence of submicroscopic infection in Africa. Historical estimates of malaria prevalence^32^ were used to assign surveys in Africa to one of three transmission intensity archetypes, and then separate models fitted to each to explore whether variation in the present and historical levels of transmission in a region might influence the frequency of submicroscopic infection in endemic populations. **(A)** Map detailing the African countries for which prevalence surveys were identified, as well as their assigned transmission archetypes based on historical and current transmission intensity (high high= historically high and currently high; high low = historically high, currently low; low low = historically low and currently low). **(B)** The microscopy: PCR prevalence ratio of surveys in each transmission archetype (dots, **n = 72, 71** and **26** for high high, high low and low low, respectively), and the modelled average prevalence ratio (coloured line) with 95% credible interval (shaded area). **(C)** The model fitted relationship (coloured lines) and 95% credible interval (shaded areas) for each of the transmission archetypes, with a particular focus on low prevalence (<20%) settings (inset box).

### The Influence of Age on Submicroscopic Parasite Carriage

Both increasing age (independent of exposure) and increased immunity (due to previous exposure, which also increases with age) have been linked to lower parasite densities^30,31^. Previous reviews of submicroscopic epidemiology have confirmed this, although data limitations have previously restricted analyses to only two age categories (≤ 16 and > 16 years old), with a limited number of datapoints in each (n = 28 and n = 13 respectively)^25^. Motivated by the substantial increase in data generated through the updated systematic review, we defined 3 age-based categories: infants (0-5 years old), older children (5-15 years old) and adults (>15 years old), yielding 40, 37 and 43 prevalence survey pairs. The prevalence ratio varied significantly between age groups (ANOVA, p<0.001, df=2), and was significantly lower in adults (indicating a greater proportion of submicrosopic infections) compared to young children (Tukey’s HSD, p<0.001) and older children (Tukey’s HSD, p<0.001). A similar disparity was observed between older and younger children, although this difference was not significant (Tukey’s HSD, p=0.61). We also explored whether these differences were observed consistently across the range of transmission settings (using survey PCR prevalence as a proxy for transmission intensity) present in the data **(Fig.4A)**. Fitting the log-linear regression model separately to the data for each age group highlighted that the increased prevalence ratio observed in infants and older children compared to adults was less pronounced in higher transmission settings **(Fig.4B)**. For example, in high endemic areas with 70% overall PCR prevalence, the prevalence ratio for infants was predicted to be 1.42x that of adults, but 1.92x at low endemic areas with 10% overall PCR prevalence. A similar result was observed for adults and older children, suggesting genuine differences in submicroscopic epidemiology both between age groups and across transmission settings.

**Figure 4:**
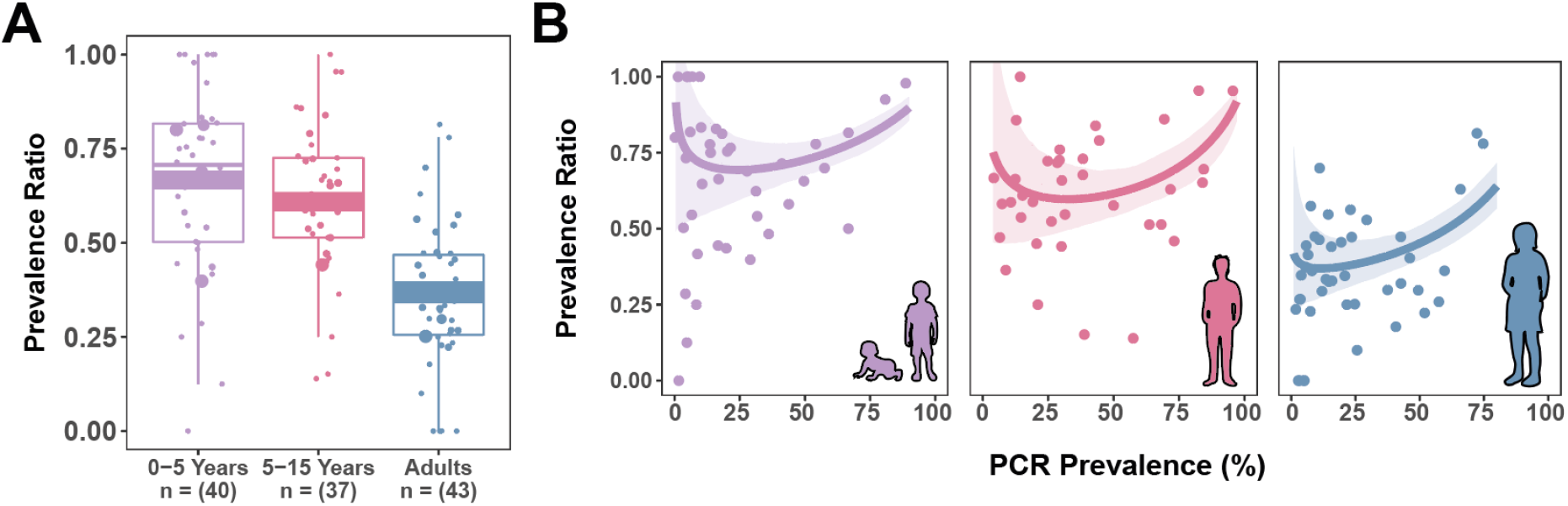
The influence of age on submicroscopic parasite carriage. Separate models were fitted where age disaggregated data were available to assess whether the prevalence of infection by microscopy compared to PCR varied with age. **(A)** Age disaggregated prevalence survey data for young children (0-5 years old, purple points, **n = 40)** older children (5-15 years old, pink points, **n = 37)** and adults (blue points, **n = 43)** with the fitted model relationship (coloured line) and 95% credible interval for each (shaded area). **(B)** The prevalence of microscopy compared to PCR (the prevalence ratio) in surveys where age-disaggregated data (dots) were available by age group, showing the fitted model relationship (coloured lines) and the 95% credible interval (shaded areas).

### Contribution of the Submicroscopic Reservoir to Onwards Transmission Across Different Settings

We next explored how the contribution of submicroscopic infections to onwards transmission might vary across settings characterised by different historical patterns of transmission. To do this, we utilised a range of estimates for the infectivity of submicroscopic infections compared to microscopically detectable infections, as well as the modelled relationship between transmission intensity and the prevalence ratio for each transmission archetype. We estimate that in transmission settings characterised by both historical and current low levels of transmission (the latter defined as <= 5% survey prevalence by PCR), submicroscopically infected individuals could account for 17.5% to 68.0% of onwards transmission. The exact figure depends on the assumption surrounding the comparative infectivity of microscopically detectable and submicroscopic infections **(Fig.5C)**. By contrast, our results suggest the contribution of the submicroscopic reservoir to transmission is less important in settings where transmission has only recently declined **(Fig.5B)** although their contribution is not irrelevant, ranging from 7.8% to 46.0% depending on the assumed comparative infectivity.

**Figure 5:**
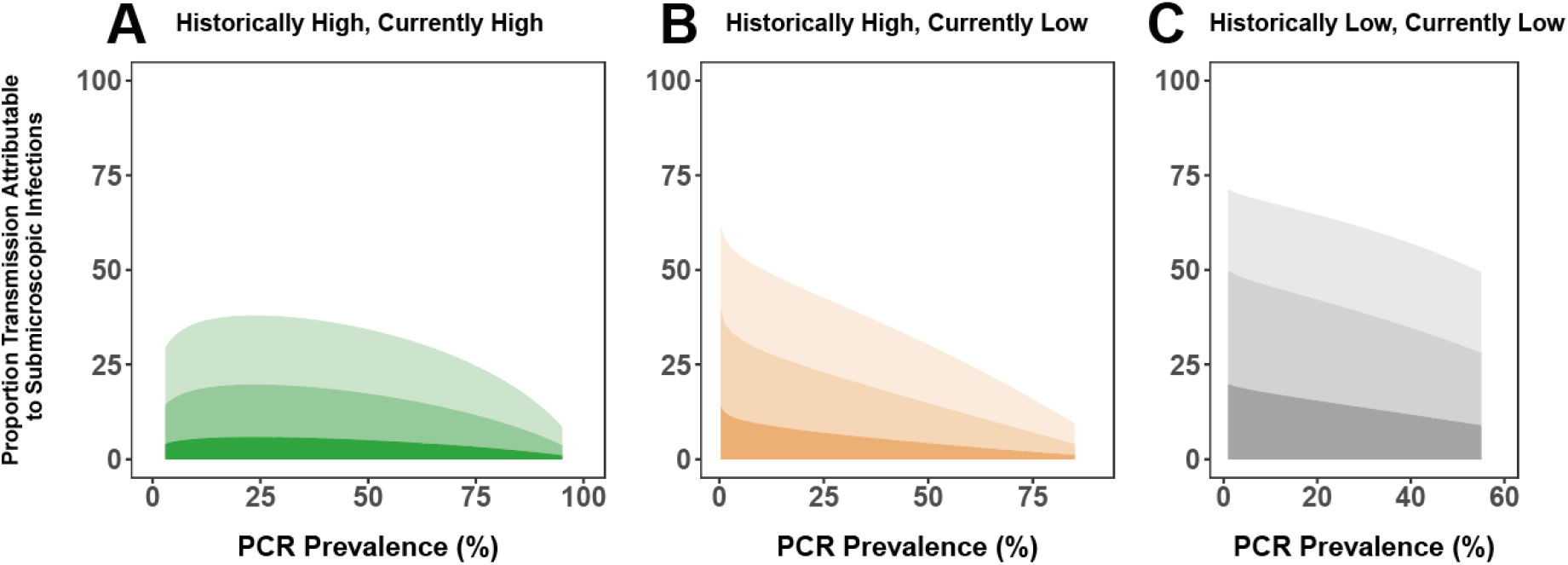
The potential contribution of submicroscopic infections to onwards transmission according to current and historical transmission intensity. Using our modelled relationship between PCR and microscopy prevalence **(Fig.3)**, we estimated the potential contribution of the submicroscopic reservoir to onwards transmission for each of the transmission archetypes **(A)** High High; **(B)** High Low; **(C)** Low Low) if microscopic infections are either 2x, 5x or 20x more infectious than submicroscopic infections. In each panel, the purple shaded area represents the proportion of transmission attributable to microscopically detectable infections, and the lower shaded areas the contribution of submicroscopic infections for each of the 3 assumed comparative infectivities-darkest shade is 20x (i.e. that microscopically detectable infections are 20x more infectious than submicroscopic infections), intermediate shade is 5x, and lightest shade is 2x.

## Discussion

Significant debate surrounds the importance of the submicroscopic reservoir to malaria control efforts and whether they need to be targeted by interventions^33^, particularly in areas of low transmission. Disaggregating the now larger quantity of available data (267 prevalence survey pairs from 38 countries) has yielded insight into the complex relationships underlying the global pattern of submicroscopic occurrence. In turn, this has facilitated a more refined evaluation of when and where submicroscopic infections are likely to be most prevalent and who is most likely to harbour them. In particular, we show that the transmission gradient in the proportion of submicroscopic infections observed across settings can be largely explained by differences in the historical patterns of transmission and the age structure of the infected population. Moreover, whilst previous work has generally noted the potential relevance of submicroscopic infections in low transmission settings^34^, our results suggest that this relevance is likely to be highly context dependent, potentially warranting different approaches to their control in different locations.

Our results highlight the importance of both individual-level factors (such as age) and setting-specific factors (such as historical patterns of transmission) in determining the size of the submicroscopic reservoir. However, an important caveat to these findings is that there were insufficient data to examine the role of these factors simultaneously. The average age of infection is typically higher in low transmission settings^35,37^; therefore, a greater proportion of infected individuals in all-age surveys would be expected to be adults. Our results surrounding past transmission history might then be confounded by differences in the average age of infection across these different settings. However, the age distribution of malaria infection appears to adapt fairly rapidly to reflect changes in transmission: in an area of south-western Senegal experiencing a 30-fold reduction in malaria incidence between 1996 and 2010, there was a shift in the age distribution of cases from predominantly under-5s to a distribution that was near uniform across the population^38^. Similarly, rapid declines in malaria between 2003 and 2007 in The Gambia were shown to be accompanied by an increase in the mean age of paediatric malaria admissions from 3.9 years to 5.6 years^39^. These clinical results also extend to the prevalence of malaria infection (as assessed by microscopy) in wider populations and communities^37^. These observations suggest that the difference in infected population age structure between surveys from historically and currently low settings and those from settings where transmission has only recently declined may not be large. Confounding due to differences in age structure are therefore unlikely to explain the observed disparity in the average size of the submicroscopic reservoir across these two types of settings.

Another potential limitation is the strong geographical bias in our transmission archetype stratification. The majority of surveys assigned to the “Historically Low, Currently Low” archetype are from East Africa, whilst the majority of surveys in the “Historically High, Currently High” category are from West Africa. The observed results across transmission archetypes could therefore in theory be reflecting geographical variation, rather than variation driven by past transmission history. Two strands of analysis refute this: firstly, that analysis of the submicroscopic reservoir across global regions revealed minimal difference in the prevalence ratio across West and East Africa. And secondly, that the proportion of infections which are submicroscopic in “Historically High, Currently Low” settings (a strata also predominantly composed of studies from East Africa) was consistently lower than that observed for “Historically Low, Currently Low” settings. Together with the results observed for South America, these observed differences support an effect of transmission history on the submicroscopic reservoir which is distinct from the age structure of the infected population and unlikely to be driven by geographical bias.

A number of hypotheses could explain these results, including systematic variation in asexual blood stage multiplication rate of *Plasmodium falciparum*^40^ or various haemoglobinopathies and human genetic traits that have been linked to lower average parasite densities^36,41^. Variation in the extent of parasite genetic diversity in settings characterised by different historical patterns of transmission and the influence of this on the breadth and rate of immunity acquisition might also contribute to the observed results. It is also not possible to definitively preclude systematic variation in diagnostic quality across settings, although this is perhaps unlikely: for example, a recent analysis found that whilst microscopy quality varies across settings, it does not do so systematically with transmission intensity^18^. Whilst our results highlight that PCR methodology does significantly influence the prevalence ratio, systematic variation in methodological quality across geographical and transmission archetypes settings was not observed. Interestingly, our analyses revealed no influence of seasonal effects on submicroscopic carriage, despite parasite densities having been shown to slightly rise during the rainy season even when prevalence does not change significantly (possibly due to increased occurrence of superinfection during these periods)^18,42^. This is consistent with other recent results which similarly found no difference in the submicroscopic reservoir between seasons, when directly comparing the wet and dry season submicroscopic prevalence across the same locations^43^.

Our work suggests that the contribution of submicroscopic infections to onwards transmission is likely to be highly variable across settings. However, this analysis is based on the detection of asexual parasites and does not provide direct insight into gametocyte densities. The relationship between asexual parasite and gametocyte density is highly non-linear^31^ and the distributions of parasite densities in the submicroscopic range can differ substantially between settings^18^. The proportion of submicroscopic infections may therefore not linearly relate to their contribution to onwards transmission. For example, whilst a recent membrane feeding study conducted in Burkina Faso and Kenya (high transmission settings) found that 45-75% of all mosquito infections were derived from submicroscopic infections^47^, only 4% of infections arose from submicroscopic individuals in a similar study carried out in Cambodia (a low transmission setting)^48^. These findings contrast with the predictions presented here and underscore the need to better resolve the relationship between submicroscopic parasite carriage, gametocyte densities, and mosquito infectivity.

Although progress has stalled recently, substantial enthusiasm still surrounds attempts to eliminate malaria^49^. Despite this however, and despite their potential relevance, our understanding of submicroscopic infections remains far from complete. Namely, do they represent a substantial source of transmission and threat to future progress? Do they need to be targeted in order to achieve malaria elimination? Whilst more work is required, our finding highlight important differences in submicroscopic epidemiology between settings and suggests the absence of a one-size-fits-all solution for malaria control targeting this infected subgroup. Such variation will likely warrant different approaches to malaria control if the infection is to be controlled most effectively in the approach to elimination.

## Acknowledgements

We thank Jenny Walldorf, Miriam Laufer, Andrea Buchwald, Lauren Cohee, Terri Taylor and Don Mathanga for kindly providing access to additional data from their original work.

## Contributions

LO and CD conceived the idea of the study. CW, LO and HS contributed to the design of the study. CW carried out the systematic review and the subsequent analyses, in consultation with LO and HS. CW and LO wrote the manuscript, with HS, TB, CD and AG all providing feedback and suggestions during manuscript drafting. All authors approved the final version of the manuscript.

## Code and Software

Complete details of the code used to analyse the collated data can be found at https://github.com/cwhittaker1000/Sub_Patent_Malaria_Analysis

## Methods

### Systematic Review Update and Data Extraction

Malaria prevalence data where both microscopy and PCR based methods had been used to determine infection were compiled, updating a previous review published in 2012^25^. Searches were carried out using PubMed and Web of Science and the search terms (“PCR” OR “Polymerase Chain Reaction”) AND “falciparum”. Records published January 2010 - July 2017 were searched, yielding 2136 records. A further two records, unpublished at the time of screening, were also included^20,47^, as were additional data collected through contacting authors of included references. Studies reporting asexual *P.falciparum* prevalence by microscopy and PCR in the same population were included. Surveys of pregnant women, where participants have been chosen on the basis of symptoms/treatment, or did not involve a population from a defined location were excluded. After screening titles and abstracts, 231 references were retained for full text evaluation, of which 60 were included. This yielded 253 new prevalence survey pairs, and 387 total when including previous reviews (which previously identified 45 relevant references)^21,25^ (**Supp Fig.1**). Submicroscopic infections were defined as those where infection was detectable by PCR but not by microscopy. The specificity of microscopy compared to PCR has been shown to be high (average 98.4%^21^); we therefore assume microscopy-positive individuals are also PCR-positive. In the small number of instances where the number of microscopically detected infections was higher than those identified by PCR (n = 9), the prevalence ratio was adjusted to 1 – this adjustment does not qualitatively alter the results described here.

### Statistical Analysis and Bayesian Linear Regression

As in previous reviews^22^, data were analysed using the regression-based methodology detailed in Sharp et al^50^:

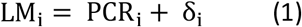

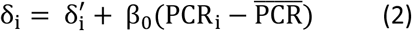

where LM_i_ = log odds of microscopy prevalence in survey *i*, PCR_i_ = log odds of PCR prevalence, 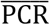 = mean survey PCR prevalence, and δ_i_ = log odds ratio (OR) of microscopy to PCR prevalence. This formulation allows δ_i_ to vary between surveys, with β_0_ controlling the extent of this variation. This model was fitted within a Bayesian Markov Chain Monte Carlo based framework, implemented in the statistical software package JAGS^51^.

### Historical and Current Regional Transmission Intensity Stratification

Surveys conducted in African settings were geolocated and prevalence estimates (aggregated to the administrative unit 1 level) from the Malaria Atlas Project^52^ used to characterise current and historical transmission intensity of the region each survey belonged to. We distinguish between local malaria transmission (defined by the prevalence recorded in each survey), and malaria transmission at the regional level (reflecting broader patterns of transmission), with this latter measure used to stratify each study into one of three “transmission archetypes”:

- **Historically High, Currently High:** areas that have historically (defined as 15 years previous to the date of the survey) high transmission intensity (>15% slide prevalence in 2-10 year olds (*Pf*PR_2-10_)) and remain so at the time of the survey (n = 71).
- **Historically High, Currently Low:** Areas of historically high transmission intensity that have declined in recent years to low levels (<15% *Pf*PR_2-10_) (n = 65).
- **Historically Low, Currently Low:** Areas characterised by historical and current low transmission (<15% *Pf*PR_2-10_) (n = 28).

Where MAP estimates were unavailable (for dates earlier than 2000), it was assumed that the year 2000 was reflective of historical transmission intensity given the only recent substantial increase in international financing for malaria control (an approximately twentyfold increase between 2000 and 2015)^53^.

### Calculating the Contribution of Submicroscopic Infections to Onwards Transmission

Estimates of comparative infectivity of microscopically-detectable vs submicroscopic infections (the “infectivity ratio”) are variable, ranging from 2x^54^ to a 20x difference^48^. We therefore explored three scenarios where microscopically detectable infections were either 2x, 5x or 20x more infectious to mosquitoes than submicroscopic infections. Proportional contribution to transmission by submicroscopic infections was calculated as:

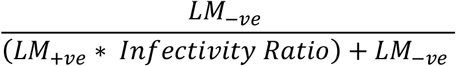

The proportional contribution to transmission is the quantity of interest: the relative infectiousness of submicroscopic infections (the prevalence of which is denoted by *LM_−ve_*) is therefore set to 1, and *Infectivity Ratio* (2x, 5x or 20x) is a multiplicative factor reflecting the fact that microscopically detectable infections (prevalence denoted by *LM_+ve_*) are more infectious. The equation’s denominator reflects total onwards transmission occurring within the population; the numerator the amount of transmission attributable to submicroscopic infections. These analyses assume that that submicroscopic and microscopically infected populations do not differ in other key factors likely to also influence transmission (e.g. age and mosquito exposure).

