## Supplementary Information for "Global & Temporal Patterns of Submicroscopic *Plasmodium falciparum* Malaria Infection"

### Contents:

**Supplementary Information Statistical Analyses**

**Supplementary Figure 1:** Systematic review overview, workflow and selection of eligible studies

**Supplementary Figure 2:** Prevalence of infection by PCR versus microscopy and model fits for previously collated data and data newly identified as part of this review

**Supplementary Figure 3:** Comparison of empirically observed and modelled microscopy prevalence

**Supplementary Figure 4:** Comparison of different model structures and their capacity to fit the collated data

**Supplementary Figure 5:** Comparing the prevalence ratio across different sampling seasons.

**Supplementary Figure 6:** Comparing the prevalence ratio across different PCR methodologies

**Supplementary Figure 7:** Tabulation of diagnostic properties by global region

**Supplementary Figure 8:** Tabulation of diagnostic properties by transmission archetype

**Supplementary Figure 9:** Sensitivity analysis to assess the robustness of the results surrounding historical and current patterns of Transmission intensity.

**Supplementary Figure 10:** Raw MCMC Output from Full Data Model Fitting

**Supplementary Figure 11:** Raw MCMC Output from Global Region Stratified Model Fitting

**Supplementary Figure 12:** Raw MCMC Output from Transmission Archetype Stratified Model Fitting

**Supplementary Figure 13:** Raw MCMC Output from Age Stratified Model Fitting

#### Supplementary Information Statistical Analyses

##### Model Formulation

The data were analysed using a meta-analytic technique detailed in Sharp et al[1], that allows for an association between LM prevalence and PCR prevalence, measured as the odds ratio, that is dependent on the underlying level of malaria transmission (as measured by PCR prevalence). Although a logit-linear model was previously used to model the relationship between LM and PCR prevalence, we initially explored a range of different model structures to assess their capacity to fit this newly collated and updated dataset. These were formulated as follows:

$$\mathrm{LM_i} = \mathrm{PCR_i} + \delta_\mathrm{i} \quad (1)$$

where:

$$\delta_\mathrm{i} = \delta_\mathrm{i}' \quad \textbf{(Basic) (2)}$$

$$\delta_\mathrm{i} = \delta_\mathrm{i}' + \beta(\mathrm{PCR_i} - \overline{\mathrm{PCR}}) \quad \textbf{(Linear) (3)}$$

$$\delta_\mathrm{i} = \delta_\mathrm{i}' + \beta(\mathrm{PCR_i} - \overline{\mathrm{PCR}}) + \gamma(\mathrm{PCR_i} - \overline{\mathrm{PCR}})^2 \quad \textbf{(Quadratic) (4)}$$

$$\delta_\mathrm{i} = \delta_\mathrm{i}' + \beta(\mathrm{PCR_i} - \overline{\mathrm{PCR}}) + \gamma(\mathrm{PCR_i} - \overline{\mathrm{PCR}})^2 + \sigma(\mathrm{PCR_i} - \overline{\mathrm{PCR}})^3 \quad \textbf{(Cubic) (5)}$$

where $\mathrm{LM_i}$ is the log odds of microscopy prevalence in trial *i* and $\mathrm{PCR_i}$ is the log odds of PCR prevalence in trial *i*. $\overline{\mathrm{PCR}}$ is the log odds of the mean PCR prevalence across surveys. $\delta_\mathrm{i}$ is the log odds ratio (OR) of microscopy to PCR prevalence, with $\delta_\mathrm{i}'$ the expected log OR when the log odds of PCR prevalence is equal to the overall mean across trials. Formulation in this way allows the $\delta_\mathrm{i}$ to vary between surveys (a phenomenon established in previous reviews[2]): $\beta$, $\gamma$ and $\sigma$ are regression coefficients specifying the extent and nature of this variation.

##### Model Fitting

Each of these models were fitted to the entirety of the collated prevalence data (including both surveys identified in this review as well as those from previous reviews[2,3]) within a Bayesian Markov Chain Monte Carlo (MCMC) based framework, implemented in the statistical software package JAGS[4]. The observed prevalence values were assumed to be drawn from a binomial distribution with the sample size of the survey as the number of trials (people tested) and the underlying true prevalence as the probability of "success" (malaria positivity) in any given trials:

$$\mathrm{Positive}_{LM,\mathrm{i}} \sim \mathrm{Binomial}(\mathrm{Prevalence}_{LM,i}, \mathrm{N})$$

$$\mathrm{Positive}_{PCR,\mathrm{i}} \sim \mathrm{Binomial}(\mathrm{Prevalence}_{PCR,i}, \mathrm{N})$$

Where $\text{Positive}_{LM,\text{i}}$ and $\text{Positive}_{PCR,\text{i}}$ are the observed number of malaria positive individuals by each diagnostic method, $\text{Prevalence}_{LM,i}$ and $\text{Prevalence}_{PCR,i}$ are the underlying "true" prevalences (and the quantity that is modelled in equations 1 – 5 above).

Uninformative prior distributions were assigned to all parameters in the model. In each instance, 4 chains of 10,000 iterations were run for purposes of model fitting and parameter inference. 5,000 of these iterations were discarded as burn-in, leaving 5,000 iterations from each chain and therefore a total of 20,000 iterations available for inference. This sample was further thinned by selecting only every 10th element in order to minimise auto-correlation, leaving a sample of 2,000 values upon which inference was based. Measures of MCMC convergence such as the Gelman-Rubin convergence statistic were monitored in all cases and were all consistently < 1.02, indicating stability of the chains and likely convergence to the underlying true posterior distribution.

#### Model Comparison

Model comparison was carried out using the deviance information criterion (DIC), which considers both the capacity of the model to fit the data, as well the model's underlying complexity[5,6]. It is formulated as follows:

$$\text{DIC} = \text{D}(\bar{\theta}) + 2p_D$$

where $\text{D}(\bar{\theta})$ is the deviance evaluated at the expectation of θ (the vector of parameter values that together specify the model used, so $\delta_{\text{i}}^{\prime}, \beta, \gamma$ and $\sigma$ for our purposes) and $p_D$ is the variance of the deviance evaluated across all values of θ. This latter quantity can be considered to be a proxy measure of the effect number of parameters the model contains, and so reflects the underlying complexity of the model. Lower DIC values are preferred and so in doing so, the DIC trades off model fit (as indicated by $\text{D}(\bar{\theta})$) and model complexity ($p_D$) to enable (in theory) selection of the model best able to extrapolate to new, unobserved data. For the data considered here, a model with a linear relationship linking PCR and microscopy prevalence on the logit scale (the model used in previous reviews) was found to have the lowest DIC (see Supplementary Figure 2), and is therefore the preferred model. Based on this, this model was used for all subsequent analyses presented in the main text of the paper.

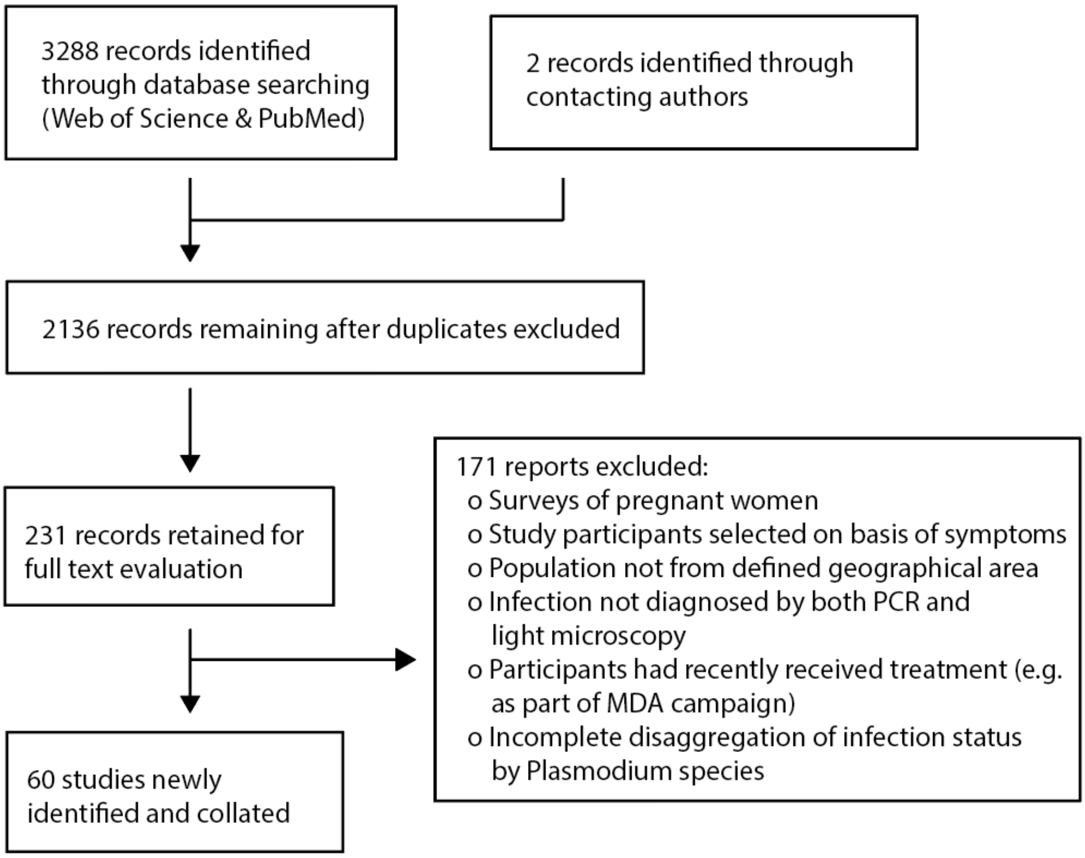


**Supplementary Figure 1: Systematic review overview, workflow and selection of eligible studies.** Searches for malaria prevalence data where infection status had been determined using both microscopy and PCR based methods. 51 studies were included in the formal analyses, including two references unpublished at the time of screening and a limited amount of additional data collected through contacting study authors.

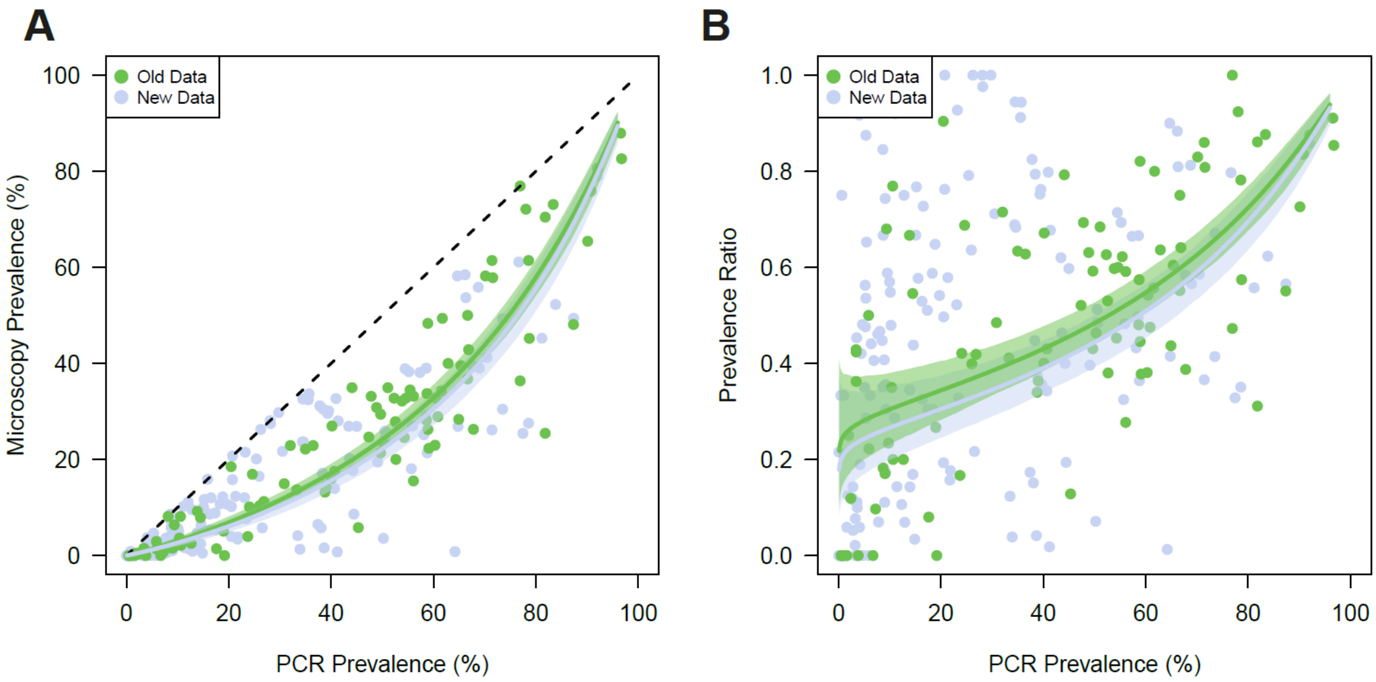


**Supplementary Figure 2: Prevalence of infection by PCR versus microscopy and model fits for previously collated data and data newly identified as part of this review.** Bayesian Markov chain Monte Carlo methods were used to fit a log-linear relationship between PCR prevalence and microscopy prevalence separately to data collated during previous reviews on submicroscopic malaria infections (n = 105, green dots) and data newly identified as part of this review (n = 177, pale purple dots) . **(A)** Microscopy and PCR Prevalence data from surveys, with the fitted model relationship (green and pale purple lines) and the 95% credible interval of the mean (shaded areas around each line). **(B)** The sensitivity of microscopy (defined here as the proportion of PCR positive individuals also detectable by microscopy) according to underlying PCR prevalence for each of the survey microscopy-PCR pairs used to fit the full model. For each dataset, the estimated average sensitivity (coloured line) and 95% credible interval of the mean (shaded area) also shown.

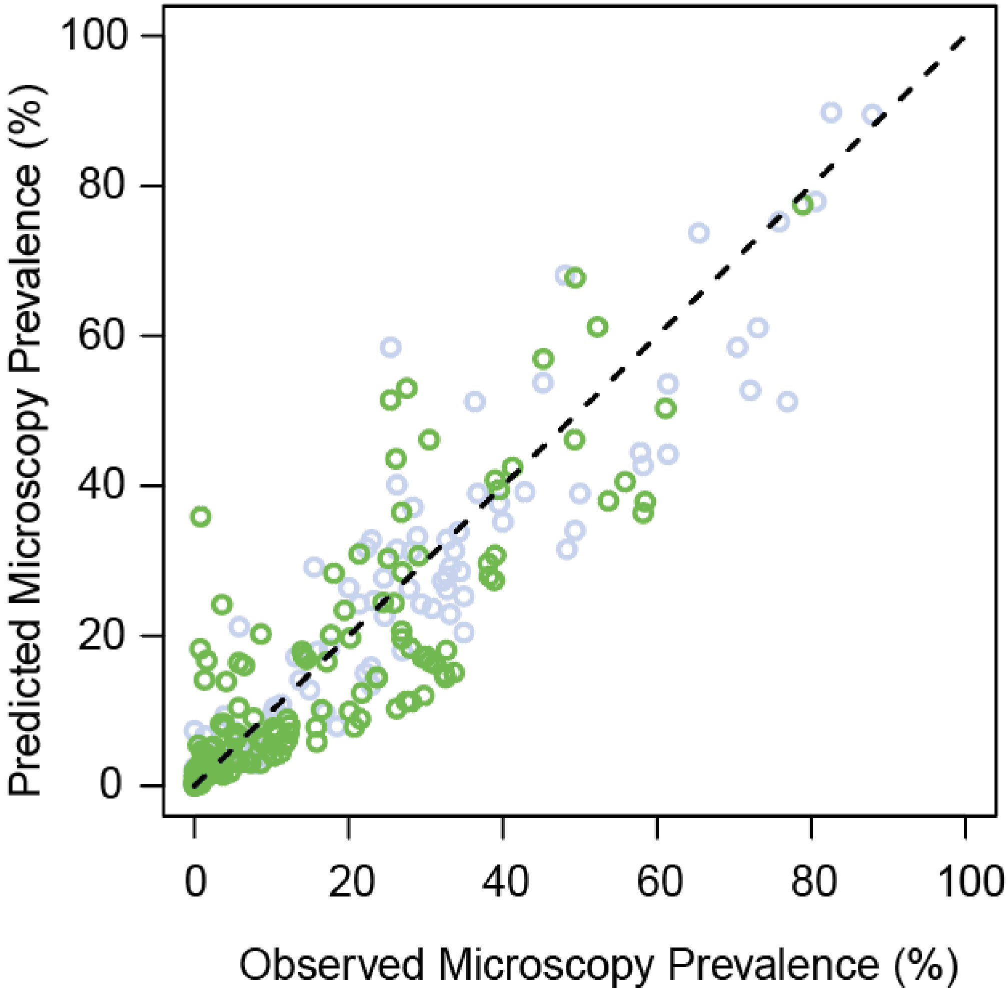


**Supplementary Figure 3: Comparison of Empirically Observed Microscopy Prevalence and Microscopy Prevalence Predicted by Bayesian Regression Modelling.** Bayesian Markov chain Monte Carlo methods were used to fit a linear relationship between PCR prevalence and microscopy prevalence on the log odds scale. For each survey (green for newly identified surveys as part of this systematic review, grey for surveys identified in previous systematic reviews), the empirically observed and modelled microscopy prevalence were plotted and compared. The correlation between the observed and model-predicted microscopy prevalence across the surveys was 0.90 (measured via $R^2$, the correlation coefficient).

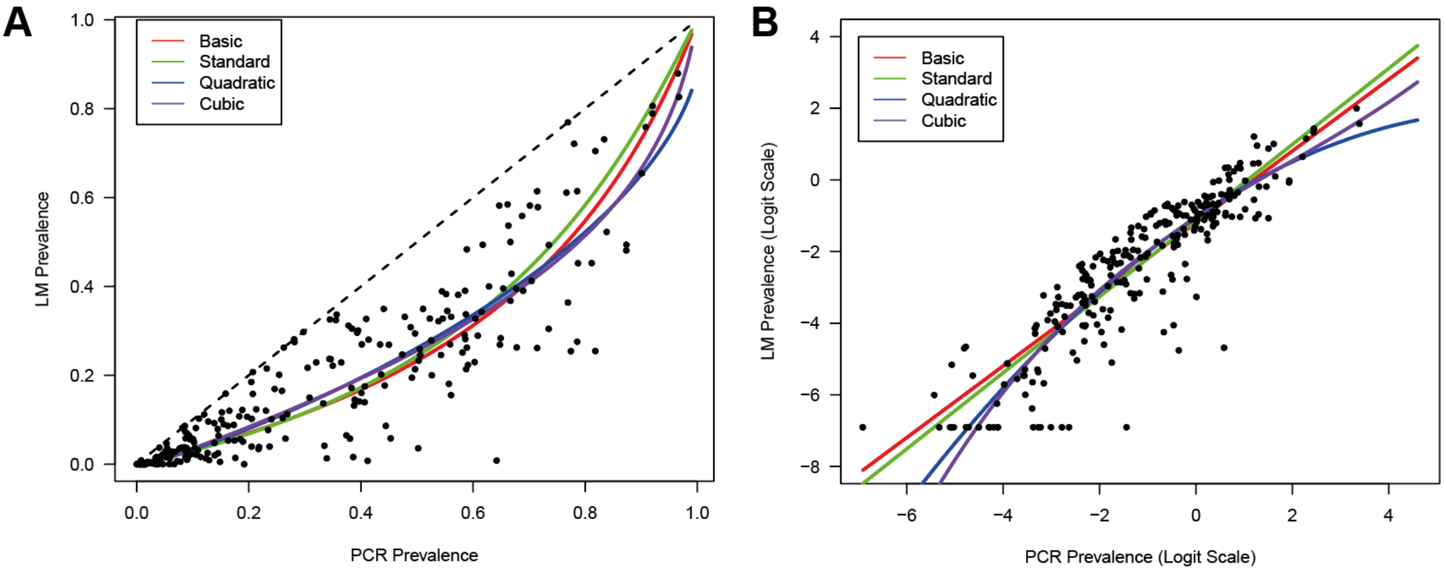


**C**

| Model Structure | Model Equation | DIC |
|---|---|---|
| Basic | $\mathrm{LM_i} = \mathrm{PCR_i} + \delta_\mathrm{i}$ | 3668.390 |
| Standard (Linear) | $\mathrm{LM_i} = \mathrm{PCR_i} + \delta_\mathrm{i} + \beta(\mathrm{PCR_i} - \overline{\mathrm{PCR}})$ | 3661.914 |
| Quadratic | $\mathrm{LM_i} = \mathrm{PCR_i} + \delta_\mathrm{i} + \beta(\mathrm{PCR_i} - \overline{\mathrm{PCR}}) + \gamma(\mathrm{PCR_i} - \overline{\mathrm{PCR}})^2$ | 3668.886 |
| Cubic | $\mathrm{LM_i} = \mathrm{PCR_i} + \delta_\mathrm{i} + \beta(\mathrm{PCR_i} - \overline{\mathrm{PCR}}) + \gamma(\mathrm{PCR_i} - \overline{\mathrm{PCR}})^2 + \sigma(\mathrm{PCR_i} - \overline{\mathrm{PCR}})^3$ | 3716.581 |

**Supplementary Figure 4: Comparison of different model structures and their capacity to fit the collated data.** Bayesian Markov chain Monte Carlo methods were used to fit a number of different relationships, varying in flexibility, to PCR and microscopy prevalence on the logit scale. **(A)** Microscopy and PCR Prevalence data from surveys (black points), with the fitted model relationship (red, green, blue and purple lines, denoting Basic, Linear, Quadratic and Cubic relationships on the logit scale respectively) all plotted on the natural scale. **(B)** As for (A), but plotted on the logit scale. **(C)** Description of the different model structures considered, along with the corresponding deviance information criterion (DIC) for each. Lower DIC values indicate a more preferred model.

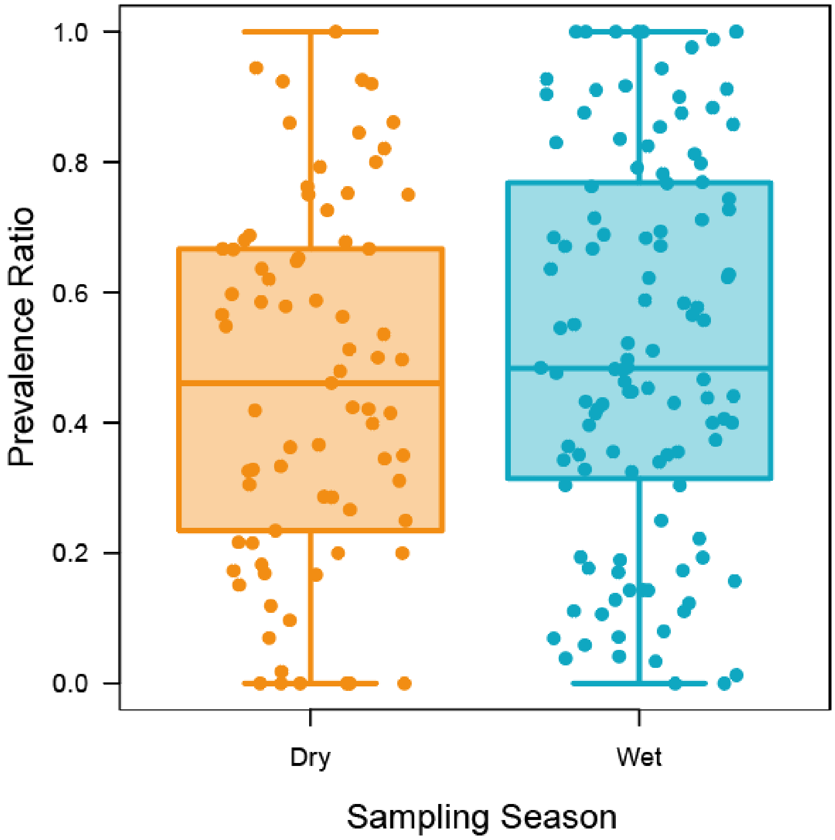


**Supplementary Figure 5: Comparing the prevalence ratio across different sampling seasons.** Where available, information from references on the seasonal timing of the sampling was extracted and collated. Presented are boxplots of the prevalence ratio (defined as the ratio of microscopically detectable infections and PCR detectable infections, with a lower prevalence ratio indicating a higher proportion of individuals with submicroscopic infections) stratified by sampling season (n = 77 for dry season sampling, and n = 116 for wet season sampling), including also the raw datapoints (coloured circles).

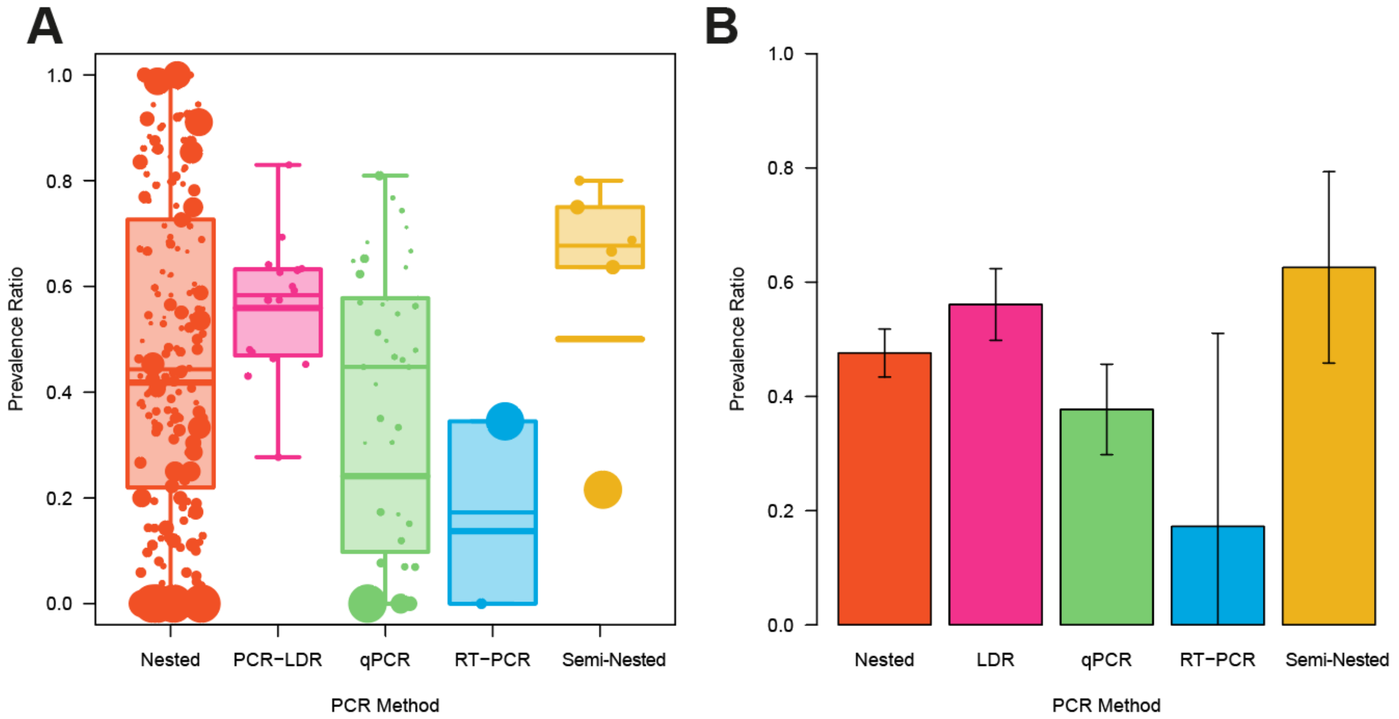


**Supplementary Figure 6: Comparing the prevalence ratio across different PCR methodologies.** Where available, information from references on the seasonal timing of the sampling was extracted and collated. **(A)** Boxplots of the prevalence ratio (defined as the ratio of microscopically detectable infections and PCR detectable infections, with a lower prevalence ratio therefore indicating a higher proportion of individuals with submicroscopic infections) stratified by PCR methodology used to determine malaria infection in the survey, including also the raw datapoints (coloured circles, weighted by the inverse of their variance), and the weighted mean (thicker horizontal lines). There was a statistically significant difference in the mean prevalence ratio across PCR methodologies ($p < 0.001$), with the prevalence ratio consistently lower in surveys using qPCR to determine infection status (indicating that microscopy performs more poorly compared to qPCR than with other PCR methodologies). This significance remained even after explicitly accounting for PCR Prevalence in the underlying model.

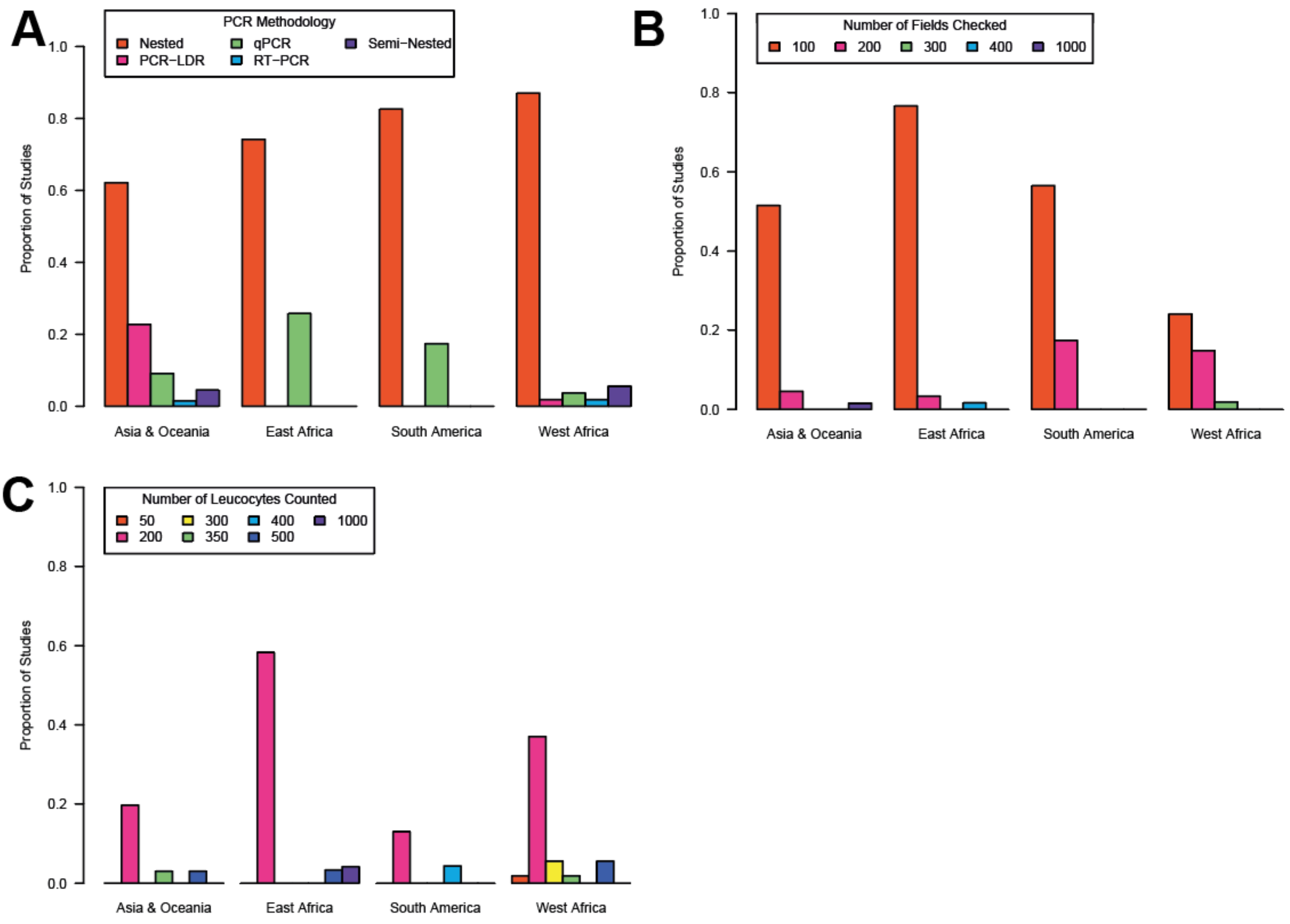


**Supplementary Figure 7: Tabulation of diagnostic properties by global region.** Where available, information from each of the references were collated detailing some of the properties of the microscopy and PCR diagnostic methodologies used to determine infection status, and then the proportion of studies using each methodology disaggregated by global region. Specifically, these were **(A)** the type of PCR used to diagnose malaria infection, **(B)** the number microscopy fields checked when examining blood smears and **(C)** the number of leucocytes counted. Note that in some instances, the number of leucocytes counted was not used to determine infection status, but instead used to calculate parasite densities. The exact purpose of leucocyte counting was not consistently reported however, and so all instances in references that mention the number of leucocytes counted are tabulated here.

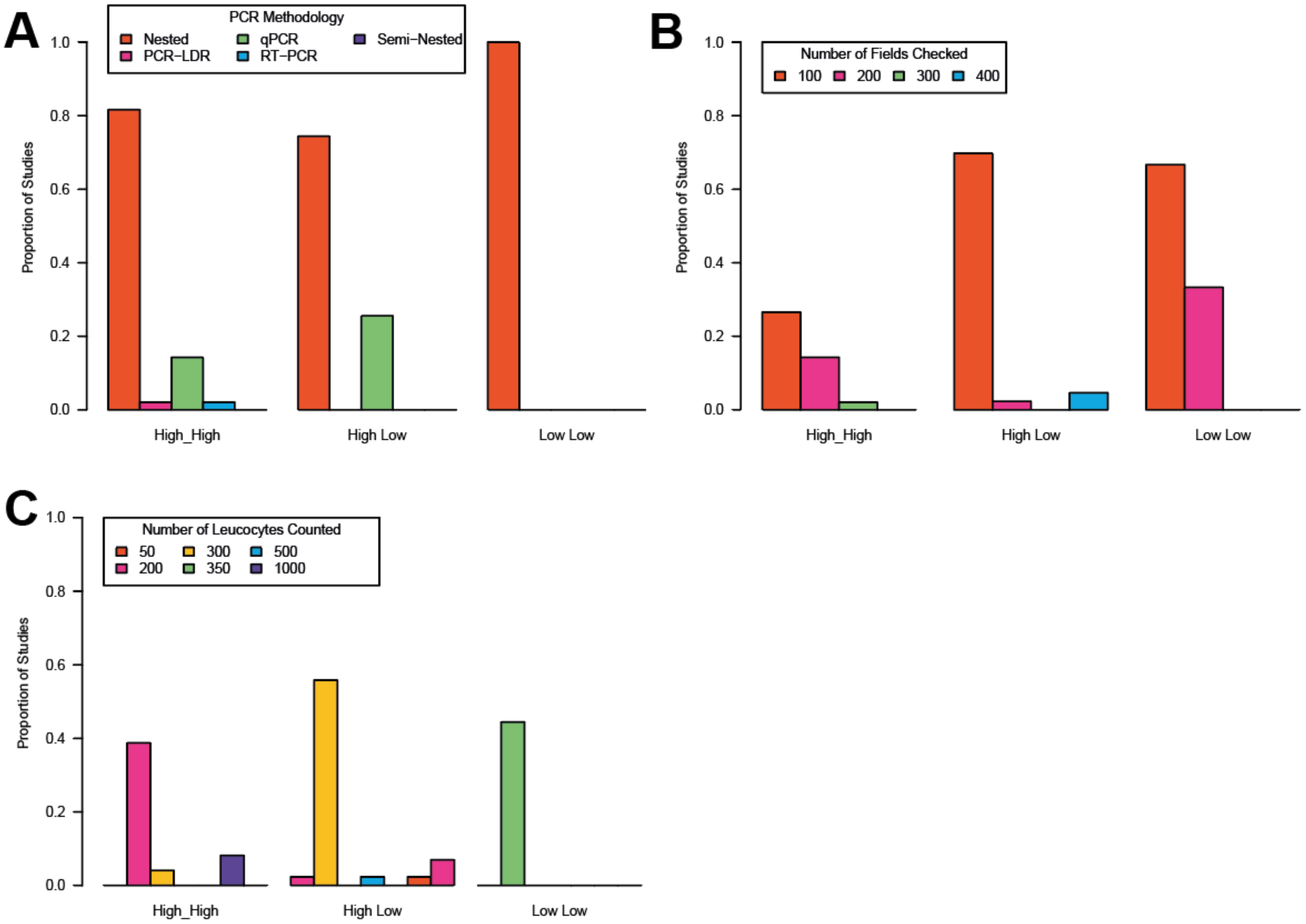


**Supplementary Figure 8: Tabulation of diagnostic properties by transmission archetype.** Where available, information from each of the references were collated detailing some of the properties of the microscopy and PCR diagnostic methodologies used to determine infection status, and then the proportion of studies using each methodology disaggregated by transmission archetype (defined as described in the Methods section of the main text). Specifically, these were **(A)** the type of PCR used to diagnose malaria infection, **(B)** the number microscopy fields checked when examining blood smears and **(C)** the number of leucocytes counted. Note that in some instances, the number of leucocytes counted was not used to determine infection status, but instead used to calculate parasite densities. The exact purpose of leucocyte counting was not consistently reported however, and so all instances in references that mention the number of leucocytes counted are tabulated here.

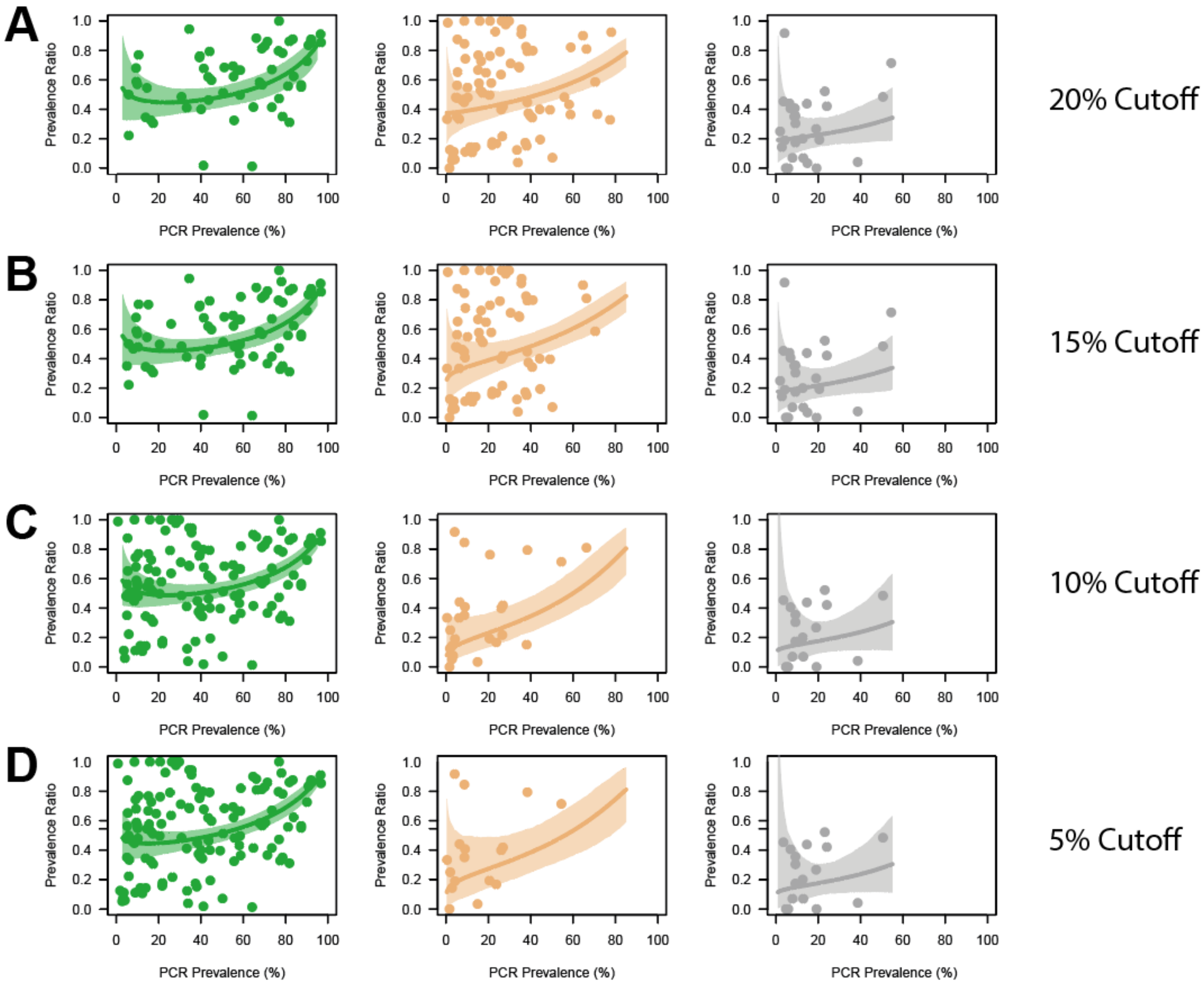


# E

| | | Comparison | | |
|---|---|---|---|---|
| **Prevalence Cutoff** | **Overall Significance** | **High High vs High Low** | **High Low vs Low Low** | **High High vs Low Low** |
| **20%** | Yes (p < 0.001) | Yes (p < 0.001) | Yes (p = 0.003) | Yes (p < 0.001) |
| **15%** | Yes (p < 0.001) | Yes (p = 0.004) | Yes (p = 0.003) | Yes (p < 0.001) |
| **10%** | Yes (p < 0.001) | Yes (p < 0.001) | No (p = 0.942) | Yes (p < 0.001) |
| **5%** | Yes (p < 0.001) | Yes (p < 0.001) | No (p = 0.914) | Yes (p < 0.001) |

**Supplementary Figure 9: Sensitivity analysis to assess the robustness of the results surrounding historical and current patterns of transmission intensity.** The analyses presented in Figure 5 of the main text Results section were repeated using different thresholds (5, 10, 15 or 20%) for defining transmission archetypes. In each instance, the standard log-linear model was fitted and the results plotted for each definition (panels **(A)** – **(D)**), where the points represent a single survey, the line represents the model fit and the pale shaded area the 95% Credible Interval. Statistical analyses were also carried out **(E)**, with ANOVA used to explore whether the mean prevalence ratio of the three transmission archetypes significantly differed (table 2$^{nd}$ column), with a post-hoc Tukey test carried out to assess pairwise differences (table columns 3-5).

**Supplementary Figure 9 Additional Text:**

In order to assess whether the threshold used to define transmission archetypes influenced our results, we repeated the analyses presented in Figure 5 of the main text (which uses a prevalence of 15% to distinguish "high" from "low" transmission) with a variety of different thresholds (specifically, 5, 10 and 20%). Surveys from sub-Saharan African countries were classified into one of three transmission archetypes (described further in the Materials and Methods section of the main text) based on Malaria Atlas Project[7] estimates for the prevalence at the administrative unit 1 level.

As detailed in the main text, it is important to distinguish between the measured survey PCR prevalence (which describes the prevalence of malaria infection in a small geographical area), and the broader regional patterns of malaria transmission (captured/described by the Malaria Atlas Project estimates which are available at the administrative unit 1 level). In this context then, surveys can belong to regions with currently "Low" transmission, but have recorded high levels of malaria prevalence in the specific area surveyed (and vice versa).

Separate models were then fitted to each of these datasets (comprising surveys carried out in regions of historically and currently high prevalence (High High), regions where transmission has historically been high but have recently declined (High Low), and regions where transmission has historically been low and remains so (Low Low). In all instances **(Supplementary Figure 9, A – D)**, model fitting revealed an incremental decline in prevalence ratio (indicating a higher proportion of individuals who were submicroscopically infected) going from High High surveys to High Low and then Low Low surveys. This was most evident with the higher cutoffs of 20% or 15% **(Supplementary Figure 9A and 9B)** – as the cutoff threshold was lowered, an increasing number of surveys that had previously been classified as "High Low" were reclassified to "High High" and the difference in prevalence ratio between "High Low" and "Low Low" settings declined. Importantly however, the difference between "High High" and "Low Low" settings remained irrespective of the definitional threshold used, a finding that is also corroborated through the results of additional statistical analyses carried out **(Supplementary Figure 9E)**.

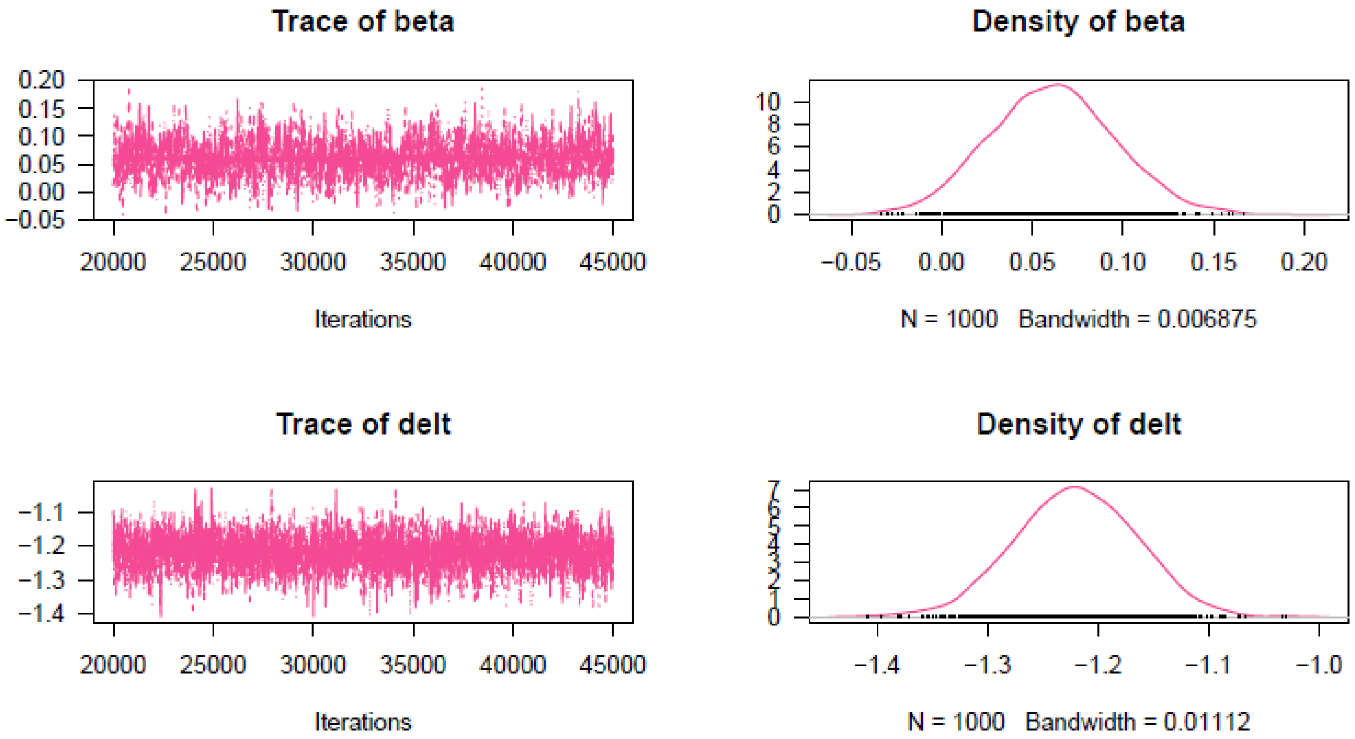


| | Mean | SD | 2.5% | Median | 97.5% |
|---|---|---|---|---|---|
| beta | 0.061 | 0.035 | -0.004 | 0.061 | 0.130 |
| delt | -1.220 | 0.055 | -1.328 | -1.220 | -1.114 |

**Supplementary Figure 10:** Raw MCMC Output from Full Data Model Fitting. A logit-linear model was fitted to age-stratified data collated during the review process. This was done within a Bayesian framework, using MCMC based inference and the statistical software package JAGS. The raw output from the MCMC sampling are presented here including the MCMC chain traces, the empirical posterior distribution, and summary statistics for each parameter including the mean, standard deviation (SD) and distribution quantiles.

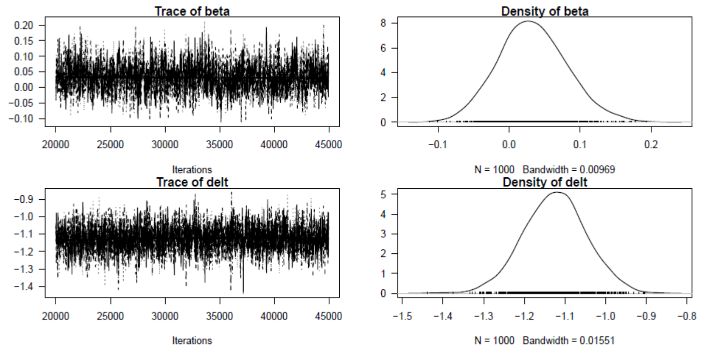


| | Mean | SD | 2.5% | Median | 97.5% |
|---|---|---|---|---|---|
| beta | 0.033 | 0.048 | -0.058 | 0.031 | 0.134 |
| delt | -1.125 | 0.077 | -1.281 | -1.124 | -0.975 |

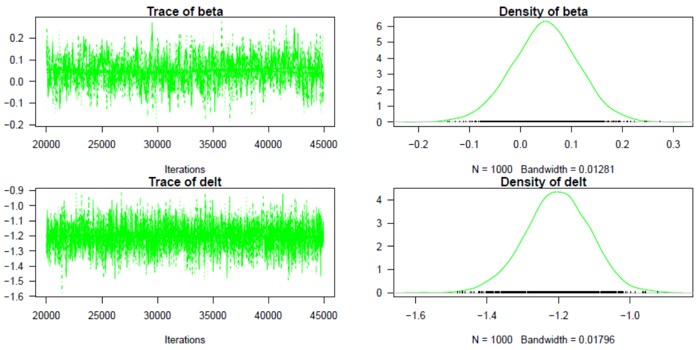


| | Mean | SD | 2.5% | Median | 97.5% |
|---|---|---|---|---|---|
| beta | 0.049 | 0.065 | -0.082 | 0.049 | 0.177 |
| delt | -1.203 | 0.090 | -1.387 | -1.202 | -1.030 |

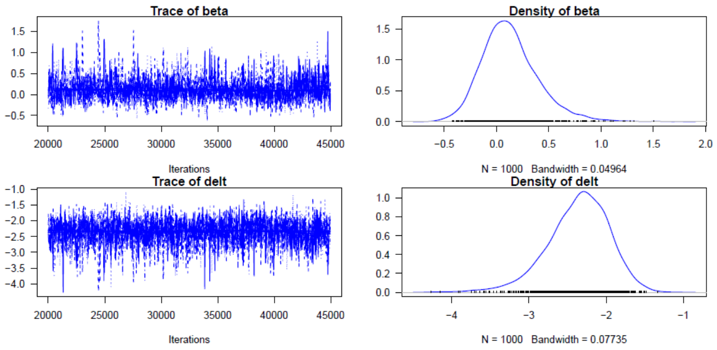


| | Mean | SD | 2.5% | Median | 97.5% |
|---|---|---|---|---|---|
| beta | 0.129 | 0.274 | -0.328 | 0.101 | 0.773 |
| delt | -2.362 | 0.403 | -3.264 | -2.325 | -1.675 |

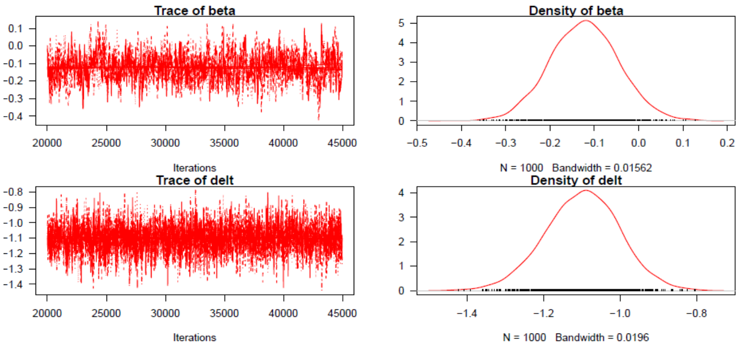


| | Mean | SD | 2.5% | Median | 97.5% |
|---|---|---|---|---|---|
| beta | -0.122 | 0.077 | -0.272 | -0.122 | 0.031 |
| delt | -1.099 | 0.097 | -1.294 | -1.096 | -0.908 |

**Supplementary Figure 11:** Raw MCMC Output from Global Region Stratified Model Fitting. A logit-linear model was fitted to global region-stratified data collated during the review process. This was done within a Bayesian framework, using MCMC based inference and the statistical software package JAGS. The raw output from the MCMC sampling are presented here including the MCMC chain traces, the empirical posterior distribution, and summary statistics for each parameter including the mean, standard deviation (SD) and distribution quantiles. Black refers to surveys carried out in Asia and Oceania, green to those carried out in East Africa, blue in South America and red in West Africa.

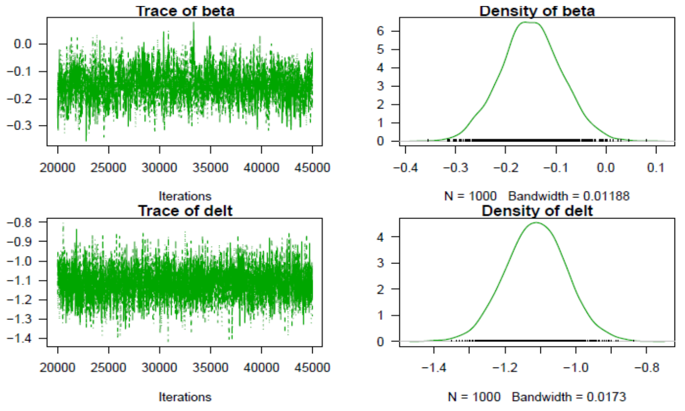


| | Mean | SD | 2.5% | Median | 97.5% |
|---|---|---|---|---|---|
| beta | 0.065 | 0.097 | -0.124 | 0.065 | 0.256 |
| delt | -1.026 | 0.113 | -1.255 | -1.023 | -0.818 |

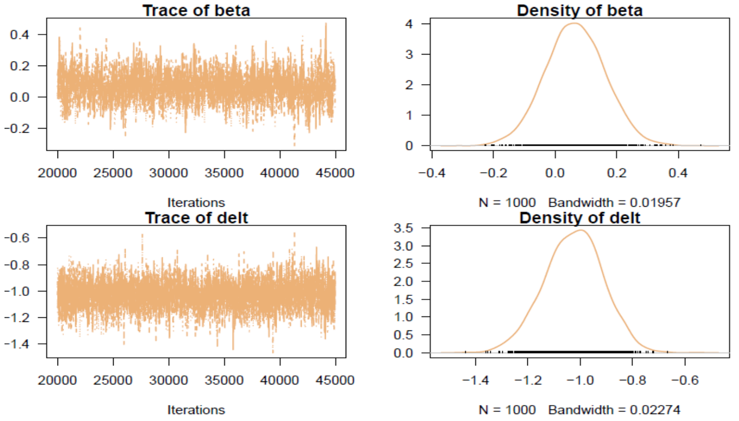


| | Mean | SD | 2.5% | Median | 97.5% |
|---|---|---|---|---|---|
| beta | 0.065 | 0.097 | -0.124 | 0.065 | 0.256 |
| delt | -1.026 | 0.113 | -1.255 | -1.023 | -0.818 |

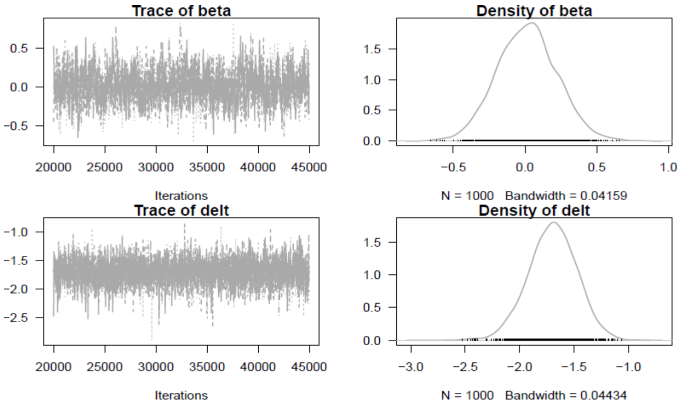


| | Mean | SD | 2.5% | Median | 97.5% |
|---|---|---|---|---|---|
| beta | 0.015 | 0.210 | -0.388 | 0.016 | 0.429 |
| delt | -1.699 | 0.225 | -2.146 | -1.692 | -1.271 |

**Supplementary Figure 12:** Raw MCMC Output from Transmission Archetype Stratified Model Fitting. A logit-linear model was fitted to transmission archetype stratified data collated during the review process. This was done within a Bayesian framework, using MCMC based inference and the statistical software package JAGS. The raw output from the MCMC sampling are presented here including the MCMC chain traces, the empirical posterior distribution, and summary statistics for each parameter including the mean, standard deviation (SD) and distribution quantiles. Green refers to surveys carried out in regions of historically and currently high transmission, tan refers to surveys carried out in areas where transmission has historically been high but has recently declined, and grey refers to surveys carried out in areas of historically and currently low transmission.

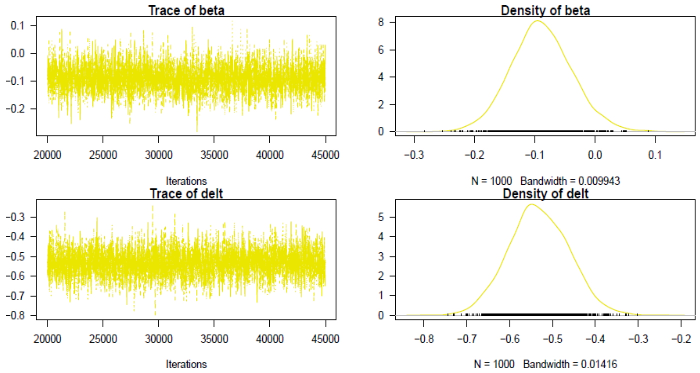


| | Mean | SD | 2.5% | Median | 97.5% |
|---|---|---|---|---|---|
| beta | -0.089 | 0.051 | -0.186 | -0.090 | 0.015 |
| delt | -0.532 | 0.070 | -0.667 | -0.534 | -0.395 |

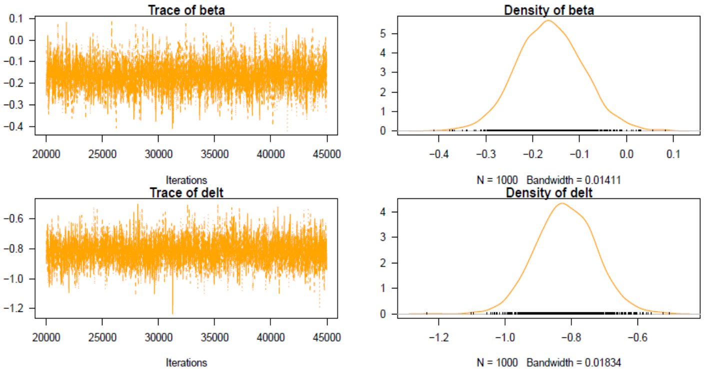


| | Mean | SD | 2.5% | Median | 97.5% |
|---|---|---|---|---|---|
| beta | -0.164 | 0.071 | -0.300 | -0.165 | -0.021 |
| delt | -0.817 | 0.091 | -0.990 | -0.818 | -0.636 |

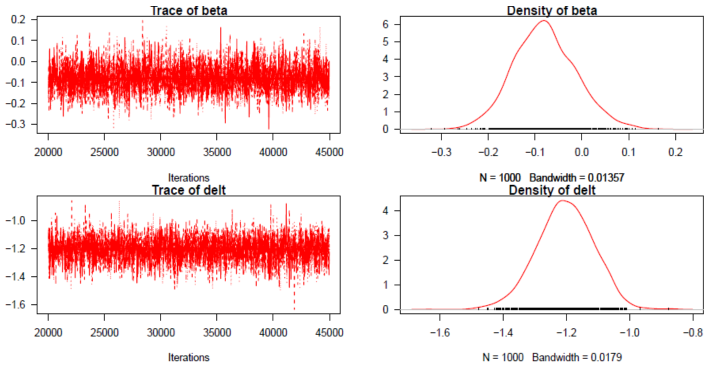


| | Mean | SD | 2.5% | Median | 97.5% |
|---|---|---|---|---|---|
| beta | -0.079 | 0.067 | -0.207 | -0.082 | 0.058 |
| delt | -1.204 | 0.090 | -1.386 | -1.203 | -1.040 |

**Supplementary Figure 11:** Raw MCMC Output from Age Stratified Model Fitting. A logit-linear model was fitted to age-stratified data collated during the review process. This was done within a Bayesian framework, using MCMC based inference and the statistical software package JAGS. The raw output from the MCMC sampling are presented here including the MCMC chain traces, the empirical posterior distribution, and summary statistics for each parameter including the mean, standard deviation (SD) and distribution quantiles. Yellow refers to young children, orange refers to older children and red refers to adults.
